# Structural and Biochemical Analysis of the CABIT1 Domain of THEMIS

**DOI:** 10.64898/2026.06.24.734275

**Authors:** Kristos I. Negrón Terón, Daniela Ortiz Salazar, Tyler S. Beyett

## Abstract

T cells are important components of the adaptive immune system and develop through a selection process regulated by signaling through the T-cell receptor (TCR). Thymocyte-Expressed Molecule Expressed in Selection (THEMIS) is a TCR-proximal protein that modulates the activity of Shp1 phosphatase to influence TCR signaling during development. THEMIS has been shown to both activate and inhibit Shp1, but the molecular mechanisms of these functions are poorly understood. THEMIS contains two rare Cysteine All-Beta In THEMIS (CABIT) domains, the N-terminal of which interacts with Shp1 and is likely responsible for modulation of its phosphatase activity. Herein, we report the first crystal structure of the THEMIS CABIT1 domain. While a portion of the CABIT1 domain is poorly resolved, it appears to share the same overall fold observed in our recent CABIT2 crystal structure and AlphaFold predictions. We show that phosphorylation of the CABIT1 domain by LCK is required for association with SHP1 and that phosphorylated CABIT1 can protect the phosphatase domain of Shp1 from oxidation and inhibition by reactive oxygen species (ROS), which may serve as a mechanism by which THEMIS enhances Shp1 activity.

## Introduction

The adaptive immune system is a key part of our response to pathogens and has been effectively leveraged in lifesaving cancer immunotherapies. Cytotoxic (CD8^+^) and helper (CD4^+^) T cells perform important adaptive immunity functions including eliminating pathogens and signaling to other immune cells, respectively. These specialized T cell subtypes develop through a highly regulated selection process that begins with double-negative thymocytes not expressing either the CD4 or CD8 co-receptors. These cells undergo positive selection in the thymus, forming double positive thymocytes expressing both the CD4 and CD8 T cell receptor (TCR) co-receptors before undergoing negative selection to yield single positive thymocytes expressing either CD4 or CD8^1^. Thymocyte-Expressed Molecule Involved in Selection (THEMIS) regulates positive selection during T cell development, and knockout of the gene leads to a significant reduction of CD4^+^ T cells, and to a lesser degree, CD8^+^ T cells^2–6^. THEMIS interacts with and modulates the activity of the tyrosine phosphatase Shp1 to modulate TCR signaling. THEMIS simultaneously binds Grb2, an adaptor protein that associates with the transmembrane protein linker of activated T cells (LAT) to localize THEMIS:Shp1 to activated TCRs^2–8^. Modulation of TCR signaling by the THEMIS:Shp1:Grb2 complex influences T cell development^9^.

The interactions between THEMIS and its binding partners are proposed to be mediated by its two CABIT domains (Cysteine-containing, all-β in THEMIS). CABIT domains are a rare type of protein domain found in four human proteins, THEMIS, Grb2-associated and regulator of Erk/MapK (GAREM), and their respective paralogs THEMIS2 and GAREM2^2, 10^. All CABIT domain-containing proteins associate with and modulate Shp family phosphatases, suggesting this as the putative function of CABIT domains^10^. How CABIT domains interact with binding partners and the consequences of these protein-protein interaction (PPIs) is current focus of the field, as the first structures of CABIT domains were only recently reported^11–12^. The first structure of a CABIT domain was of the crystal structure of the CABIT2 domain of THEMIS, which revealed CABIT domains adopt a novel domain fold^11^. Following this single-domain structure, a cryo-electron microscopy (cryo-EM) structure of Grb2 bound to the THEMIS CABIT domain tandem revealed how Grb2 recognizes a C-terminal proline-rich sequence following the CABIT2 domain of THEMIS. Determination of this structure required use of an antibody fragment that bound the CABIT2 domain, which may have introduced structural artifacts^12^.

No structure of THEMIS in complex with a phosphatase has been reported. However, a recent study showed that phosphorylation of Y34 in the CABIT1 domain of THEMIS by lymphocyte-specific protein tyrosine kinase (LCK) promotes association with and activation of Shp1 to attenuate TCR pathway signaling^13^. It was proposed that activation of Shp1 by phosphorylated THEMIS occurs through a “substrate priming” mechanism in which THEMIS disrupts the autoinhibited conformation of Shp1 to drive the phosphatase conformation into the open, active confirmation^13^. While this mechanism aligns with observations that THEMIS activates Shp1, other reports show THEMIS can inhibit Shp1 via reactive oxygen species (ROS), which oxidize the catalytic Cys^14^. These opposing functions likely occur in a condition-dependent manner and enable THEMIS to achieve required TCR signaling thresholds during development and respond to large changes in ROS like those observed during chronic viral infection^15^.

Despite these recent structural and mechanistic advances, the details of how ROS affect Shp1 in the presence of THEMIS remain unclear. We hypothesized that since phosphorylated THEMIS was reported to be a substrate of Shp1^13^, phosphorylated THEMIS would protect Shp1 from oxidation and inhibition in a substrate-competitive manner. This would represent a new mechanism by which THEMIS enhances the activity of Shp1. To better understand the structure and function of the THEMIS CABIT1 domain, we identified constructs suitable for crystallization and determined a 3.8 Å crystal structure of the CABIT1 domain. We report that CABIT1, and to a greater extent phosphorylated CABIT1, attenuates inhibition of Shp1 by ROS to preserve enzymatic activity. These data reveal a mechanism by which THEMIS maintains Shp1 activity, which may enable THEMIS to attenuate TCR signaling under mild oxidizing conditions.

## Method

### Antibodies

Phospho-tyrosine (4G10) mouse monoclonal antibody (mAB) (Cell Signaling Technology #96215S), 6x-His-tag mAB (HIS.H8) Dylight 680 (Invitrogen #MA1-21315-D680), and IRDye 800CW goat anti-mouse IgG2b (Li-Cor #926-32352) were purchased from commercial vendors and diluted according to manufacturers’ recommendations for western blotting.

### Expression and purification

All protein constructs were cloned or mutated using Q5 High-Fidelity DNA Polymerase (New England Biolabs) and verified via whole-plasmid sequencing (Plasmidsaurus). Human THEMIS CABIT1 domain constructs were cloned into pMCSG7 with an N-terminal 6xHis tag, pMCSG9 vector with an N-terminal 6xHis-maltose-binding protein (MBP) fusion tag, and pMCSG10 vector with an N-terminal 6xHis-glutathione S-transferase (GST) fusion tag, all followed by a Tobacco etch virus (TEV) protease cleavage site and transformed into *Escherichia coli* strain BL21(DE3) cells. Constructs of human Shp1 (*PTPN6*) phosphatase domain (243-530) were also cloned into pMCSG7 and pMCSG10 vectors and transformed into *Escherichia coli* strain BL21 (DE3) cells. A construct of the full-length human Grb2 was also cloned into pMCSG7 vector and transformed into *Escherichia coli* strain BL21 (DE3) cells. A construct of the CABIT2 domain of THEMIS (267-564) was also cloned into pMCSG7 vector and transformed into *Escherichia coli* strain BL21 (DE3) cells. All proteins were expressed in *E. coli* BL21(DE3) cells in Luria Broth (LB) medium at 37 °C to an optical density at 600 nm (OD600) of 0.6-0.8, cooled to 18 °C, and induced overnight with the addition of 0.2 mM isopropyl β-D-1-thiogalactopyranoside (IPTG). Pelleted cells were resuspended in buffer comprised of 50 mM Tris pH 8.0, 200 mM NaCl, 1 mM tris(2-carboxyethyl) phosphine (TCEP), and 5% glycerol (equilibration buffer). The resuspended cells were lysed through sonication. The lysate was centrifuged for 20 minutes at 70,000 g at 4 °C, and the supernatant was passed through a column of 2 mL Ni–NTA resin pre-equilibrated with equilibration buffer. The resin was washed with 25 column volumes of equilibration buffer supplemented with 40 mM imidazole and eluted from the column using equilibration buffer supplemented with 200 mM imidazole. The eluted protein was further purified by S200 size-exclusion chromatography in equilibration buffer. The purity of the proteins was verified by sodium dodecyl sulfate polyacrylamide gel electrophoresis (SDS–PAGE), aliquots flash frozen in liquid nitrogen, and stored at -80 °C.

A construct of the human LCK kinase domain (245-501) was cloned into pH7pFB vector with an N-terminal, TEV-cleavable 6xHis tag and transformed into DH10Bac competent cells. Recombinant baculovirus was prepared according to the Bac-to-Bac protocol and used to infect Sf9 insect cells, which were harvested after 65-70 h. Cells were pelleted and resuspended in buffer comprised of 50 mM Tris pH 8, 200 mM NaCl, 1 mM TCEP, and 5% glycerol (equilibration buffer). The resuspended cells were lysed through sonication. The lysate was centrifuged for 1 hour at ≥200,000 g at 4 °C and the supernatant passed through a self-packed column of 2 mL Ni–NTA resin pre-equilibrated with equilibration buffer. The resin was then washed with 25 column volumes of equilibration buffer supplemented with 40 mM imidazole and eluted from the column using equilibration buffer supplemented with 200 mM imidazole. The eluted protein was further purified by S200 size-exclusion chromatography in equilibration buffer. The purity of the enzyme was verified by SDS-PAGE, aliquots flash frozen in liquid nitrogen, and stored at - 80 °C.

### Crystallography Methods

Robotic crystallization screening was performed using commercial screens in sitting drop format with 50 µL reservoir solution and drops containing 0.5 µL protein and 0.5 µL reservoir solution. Screens were incubated at room temperature. Crystals were optimized in hanging drop format with drops containing 1 µL protein at 3-4 mg/mL and 1 µL reservoir solutions over wells containing 400 µL reservoir solution composed of 50 mM Tris pH 7.0-8.0, 0.1-0.4 M MgCl_2_, 10- 20% (w/v) PEG 4,000, and 5-15% glycerol. Small (<50 µm) clusters of plate-like crystals grew in 1-4 weeks. Crystals were briefly cryoprotected in 50 mM Tris pH 8.0, 0.2 M MgCl_2_, 20% (w/v) PEG 4,000, and 12% glycerol prior to flash freezing in liquid nitrogen.

Diffraction data were collected at the Advanced Light Source (ALS) at Lawrence Berkeley National Laboratory at 100K. Data were indexed, integrated, and scaled using DIALS (v3.21.1) via Xia2 (v3.21.1)^16–18^. The CABIT1 structure was phased using an AlphaFold prediction as a model for molecular replacement using Phaser in the Phenix program suite^18–21^. Structure refinement was performed using Phenix (v.1.21.2-5419) with rounds of iterative model building in COOT (v.0.9.8.93)^22^. The structure has been deposited in the Protein Data Bank with the accession code 10WM.

### Phosphorylation of THEMIS CABIT1

Purified THEMIS CABIT1 was phosphorylated by incubating CABIT1 with 1 μM recombinant LCK in equilibration buffer supplemented with 10 mM MgCl_2_. Reactions were initiated by the addition of 1 mM ATP, allowed to proceed for 2 hours at room temperature, and quenched with the addition of 25 mM EDTA. CABIT1 was then purified via S200 size-exclusion chromatography using a buffer comprised of 50 mM Tris pH 8.0, 200 mM NaCl, 1 mM TCEP, and 5% glycerol (equilibration buffer) and phosphorylation verified via Western blot with anti-phosphotyrosine antibody 4G10.

### Immunoblotting

After samples were ran on 12% SDS PAGE gels, western blots were run on the Trans-Blot Turbo transfer system (Bio-Rad #1704150) for 10 minutes with nitrocellulose membranes (Bio-Rad #1620115). Immunoblotting was carried out with mouse 4G10 (1:5000) and developed with goat anti-mouse IgG2b antibody conjugated with an infrared dye detected at 800 nm (1:5000) or 6x-His tag conjugated with an infrared dye detected at 680 nm (1:5000) via a Licor Odyssey FC 2800 Western Blot imaging system Pred XF (Li-Cor #53131).

### Pulldown assays

Pulldown assays were performed with N-terminal GST-fusion proteins of Shp1 C453S, Shp1 PTP (residues243-530) C453S, and Shp1 PTP C453S/D419A as the prey protein for the CABIT1 domain. To detect interactions with THEMIS, 50 μL of 50% slurry of glutathione-agarose beads was equilibrated with 1 mL of 50 mM Tris pH 8.0, 200 mM NaCl, 1 mM TCEP, and 5% glycerol (equilibration buffer). CABIT1 and Shp1 were incubated at a 1:1 ratio each at a final concentration of 20 μM for 30 minutes at room temperature before equilibrated glutathione-beads were added. The beads were incubated with the proteins for 30 minutes at room temperature before being washed twice by adding 1 mL of equilibration buffer, centrifuging at 1,000 g for 1 minute, and decanted. Bound proteins were eluted by addition of SDS (Laemmli) loading buffer, heating the beads for 5 minutes at 95°C, centrifuging for 10 minutes at 15,000 g, and decanting for SDS-PAGE and Western Blot analysis.

### Phosphatase Assays

Phosphatase assays were performed utilizing 6,8-Difluoro-4-Methylumbelliferyl Phosphate (DiFMUP, Invitrogen) dissolved in dimethyl sulfoxide (DMSO) as a fluorescent substrate for phosphatase activity with varying concentrations of H_2_O_2_ prepared by serial dilutions. Purified Shp1 phosphatase domain (PTP), CABIT1, phosphorylated CABIT1, Grb2, CABIT2, and BSA Fraction V (GE Lifesciences HyClone), which was reconstituted in a buffer containing 50 mM Tris pH 8.0, 200 mM NaCl, and 5% glycerol, were used at 1 μM and diluted in buffer composed of 50 mM Tris pH 8.0, 200 mM NaCl, and 5% glycerol. Shp1 and CABIT1 were incubated together for 5 minutes at room temperature in a volume of 5 µL at 2x the final concentration in a 384-well low-binding black plate (Greiner). Reactions were initiated by adding 5 µL of a H_2_O_2_ and DiFMUP solution for a final reaction volume of 10 µL. The data were continuously read on a CLARIOstar Plus plate reader (BMG Labtech) with one measurement per minute for 30 minutes using an excitation wavelength of 358 nm and emission wavelength of 620 nm. Data were analyzed using GraphPad Prism 10 (Dotmatics). The raw data was first analyzed using one-phase association to obtain K_obs_ values with the following formula:

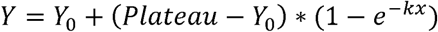

Where x is time, Y is fluorescence intensity (FI), Y_0_ is the initial FI at 0 minutes, and k is K_obs_, which can be derived to determine the inactivation efficiency, k_inact_/K_i_. The K_obs_ results were then graphed against the different concentrations of H_2_O_2_ and analyzed using non-linear and linear regression fits for the K_obs_ values observed in absence and presence of CABIT1 respectively, defined by the following formulas:

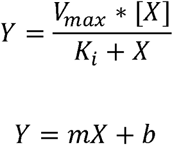

Where Y is K_obs_ and X is the H_2_O_2_ concentration. For the non-linear regression, a Michaelis-Menten fit was repurposed by taking the assumption that k_cat_/K_m_ is equal to k_inact_/K_i_ and that V_max_ referred to the point at which the phosphatase domain was saturated with H_2_O_2_ thus allowing us to solve for k_inact_ and K_i_. Assays were done in triplicate, and one-way ANOVA with Tukey post-hoc correction for multiple comparisons. were performed to determine standard deviation for each reaction and if there is a significant difference between Shp1 with or without CABIT1 and pCABIT1.

## Results

### Elucidation of the CABIT1 domain structure

CABIT1 constructs were based on domain boundaries described in previous publications, annotations in bioinformatic databases, and by inspection of the predicted AlphaFold structure of full-length THEMIS. We conducted extensive robotic screening of >1200 conditions from commercial screens using several CABIT1 constructs with and without TEV cleavage of the N-terminal His-tag. We were only able to obtain crystals in one condition using a subset of constructs with cleaved His-tags (Supplemental Table 1). The best diffracting crystals were obtained from constructs spanning residues 11-260 and 15-260 at 3-4 mg/mL using 50 mM Tris pH 8.0, 0.4 M MgCl_2_, 20% (w/v) PEG 4,000, and 12% glycerol as crystallization conditions. Despite extensive optimization and diffraction experiments on hundreds of crystals, the best crystals only diffracted to ∼3.8 Å with poor diffraction quality due to the tendency for multiple crystals to grow into one another. Here, we report the 3.8 Å crystal structure of the 15- 260 CABIT1 construct (Figure 1, Supplemental Table 2). Our previous CABIT2 structure, the first reported crystal structure of a CABIT domain, was experimentally phased using heavy atom soaks due to the anticipated novelty of the CABIT domain fold^12^. Having determined that AlphaFold accurately predicted the CABIT2 fold, we opted to use a predicted model for molecular replacement of CABIT1.

**Figure 1:**
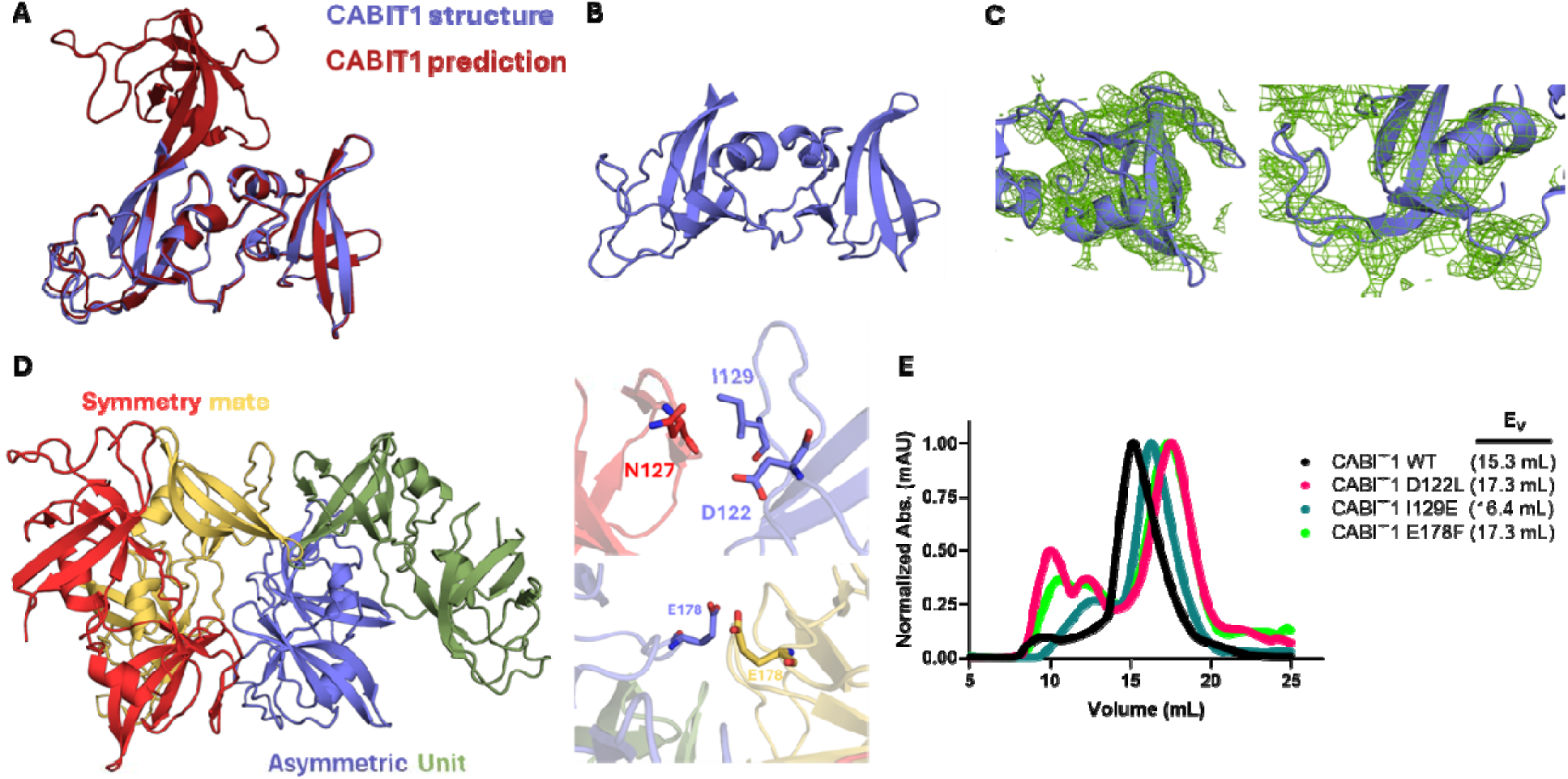
**A)** Alignment of one copy of CABIT1 found in the non-crystallographic symmetry (NCS) dimer and the AlphaFold prediction of CABIT1 (experimental structure is in slate, prediction is in brick red). **B)** One copy of the partial CABIT1 domain found in the NCS dimer. **C)** Electron density of the core subdomain and the head subdomain of the CABIT1 structure (2*F_O_- F_C_* at 1σ). **D)** Left, CABIT1 NCS dimer (slate and green) with its symmetry mate (red and wheat). Right, key contacts between symmetry mates. **E)** SEC comparison of CABIT1 WT (black) to three different CABIT1 variants (D122L: pink, I129E: teal, E178F: light green) (E_V_= elution volume).

Our CABIT1 structure aligns well with the model provided by AlphaFold, with an overall root mean square deviation (RMSD) of 0.4 Å (Figure 1A). There are two CABIT1 molecules in the asymmetric unit with chain A being better resolved. In both cases, portions of the “core” of the domain approximately spanning residues 15-54 and 231-260 are not well resolved and left unmodeled (Figure 1B), though there is sufficient space for these residues in the crystal lattice. While there is reasonable electron density for most of the protein backbone, few residues have well-resolved side chain density given the low resolution of the data (Figure 1C). We note that during purification, we observed an unexpectedly large apparent molecular weight by size exclusion chromatography and suspected a solution dimer of the isolated CABIT1 domain that is not observed for the isolated CABIT2 domain (Supplemental Figure 1). We hypothesized that the non-crystallographic dimer is the solution-state form of purified CABIT1. When analyzing the non-crystallographic symmetry (NCS) dimer, we did not observe interactions between both CABIT1 domain molecules that may facilitate dimerization. When we crystallographic symmetry mates of the NCS dimer we noted several points of contact between CABTI1 domains that could drive dimerization (Figure 1D). To test this, we selected residues at the dimer interface of NCS dimer and the crystallographic symmetry mate and mutated them with the intent to disrupt interface contacts. The D122L and E178F variants display a shift in their elution profiles via SEC towards a lower apparent molecular weight, supporting the hypothesis that the dimer observed in the crystal structure represents the solution-state form of purified CABIT1 (Figure 1E). This solution dimer is unique to the CABIT1 domain since solution dimerization is not observed when purifying isolated CABIT2 domain or THEMIS containing tandem CABIT domains (residues 1- 564) (Figure S1). Given that isolated CABIT1 domains are not found in T cells, this observation is likely an artifact of the protein constructs used for structural studies.

### Comparison to previous THEMIS structures

The crystal structure of the CABIT2 domain revealed a novel protein fold based on structural homology searches via the DALI server, which yielded no matches^11^. Similarly, our CABIT1 structure does not have structural homology to proteins in the DALI server but does contain the same two distinct sub-domains observed in the CABIT2 domain. These have been termed the “core”, consisting of several short β-sheets and two extended β-sheets all spanning through both the N- and C-terminal regions of the domain (approximately residues 1-89 and 178-260), and SH3-like “head” (approximately residues 90-177) (Figure 2A). Alignment of the CABIT1 AlphaFold prediction and CABIT2 domains revealed a large root mean square deviation (RMSD) value of 13.6 Å (Figure 2B) due to differential positioning of the head subdomain, which is nearly in line with the core in CABIT2 but rotated about 90° in CABIT1 (Figure 2C). Individual head and core subdomain alignments have RMSD values of 1.4 Å and 17.7 Å, respectively. The head subdomain is largely conserved between both domains, (Figure 2D). The largest difference with core subdomains stems from two extended, antiparallel β-sheets (β5 and β18) that are seen in CABIT2, whereas these are shorter in CABIT1. The second difference is that the SH3-like portion of the head subdomain (β10 and β11) has a longer connecting loop on CABIT 2 than in CABIT1.

**Figure 2:**
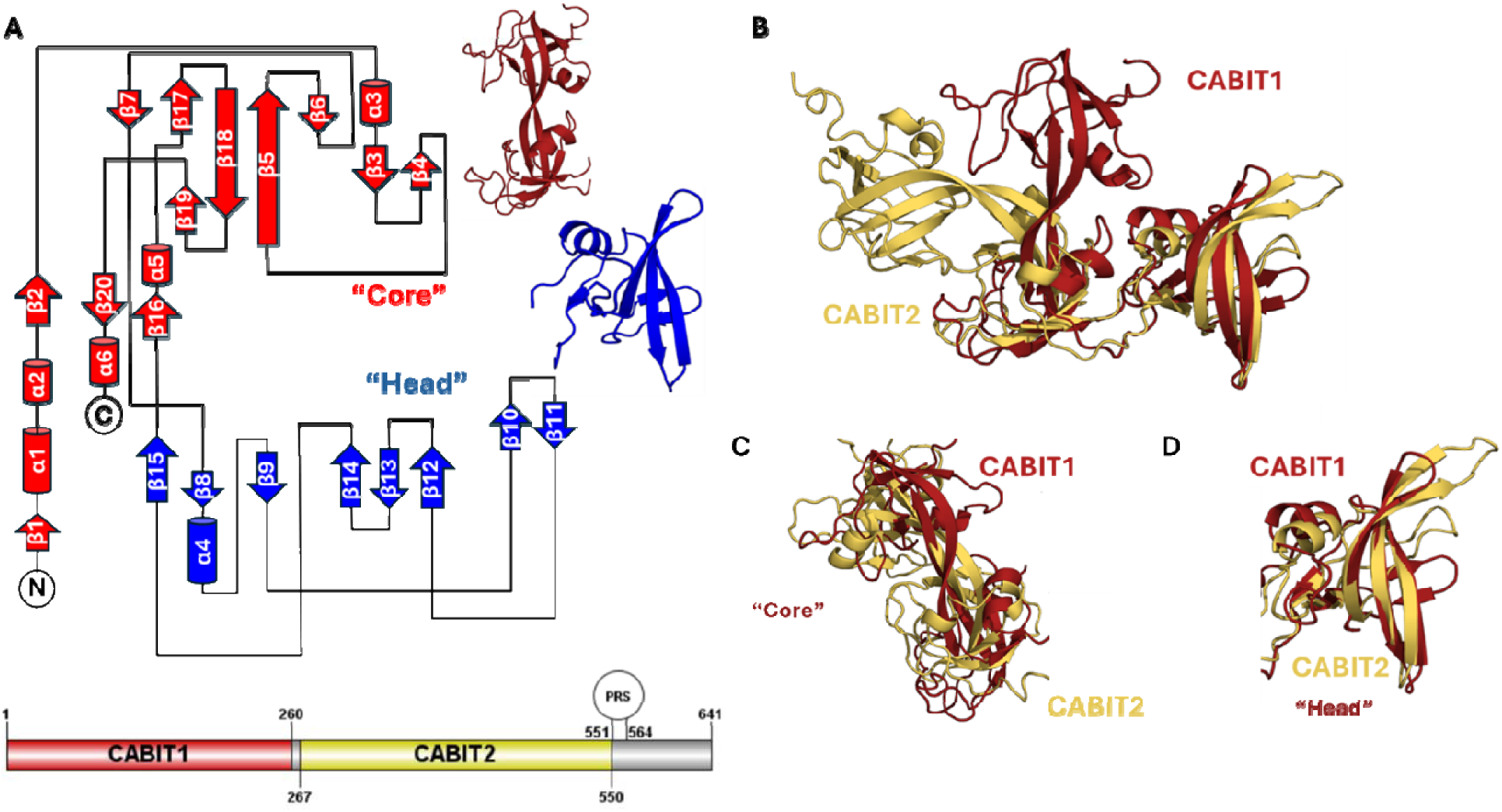
**A)** Secondary structure topology diagram of the CABIT1 domain based on AlphaFold prediction model (head subdomain: red and core subdomain: blue). **B)** Structural alignment of the CABIT1 domain (slate) and the CABIT2 domain (yellow) of THEMIS. **C)** Aligned head subdomains. **D)** Aligned core subdomains.

### Phosphorylated CABIT1 domain is required for binding to Shp1

Phosphorylation of CABIT1 at Y34 by LCK was recently shown to be required for association with and modulation of Shp1^13^. While prior experiments have shown Shp1 and THEMIS interact in a phosphorylation-dependent manner, these have only been done with either THEMIS phosphopeptides or via cellular pulldown, where we planned to analyze this using purified proteins^3,4,7,9,13^. The loop containing Y34 is unresolved in our structure, which is common for many kinase substrates. We phosphorylated CABIT1 (1-268) with purified, constitutively active LCK kinase domain to assess phosphorylation of purified CABIT1 via western blot (Figure S2). We performed GST-pulldown assays with an inactive C453S variant of the Shp1 phosphotyrosine phosphatase (PTP) also containing the secondary substrate trapping D419A mutation^24^. As a substrate, we used CABIT1 domain with and without phosphorylation by LCK to assay Shp1 binding to purified CABIT1. We observed weak association between unphosphorylated CABIT1 and robust association with phosphorylated CABIT1 (Figure 3).

**Figure 3:**
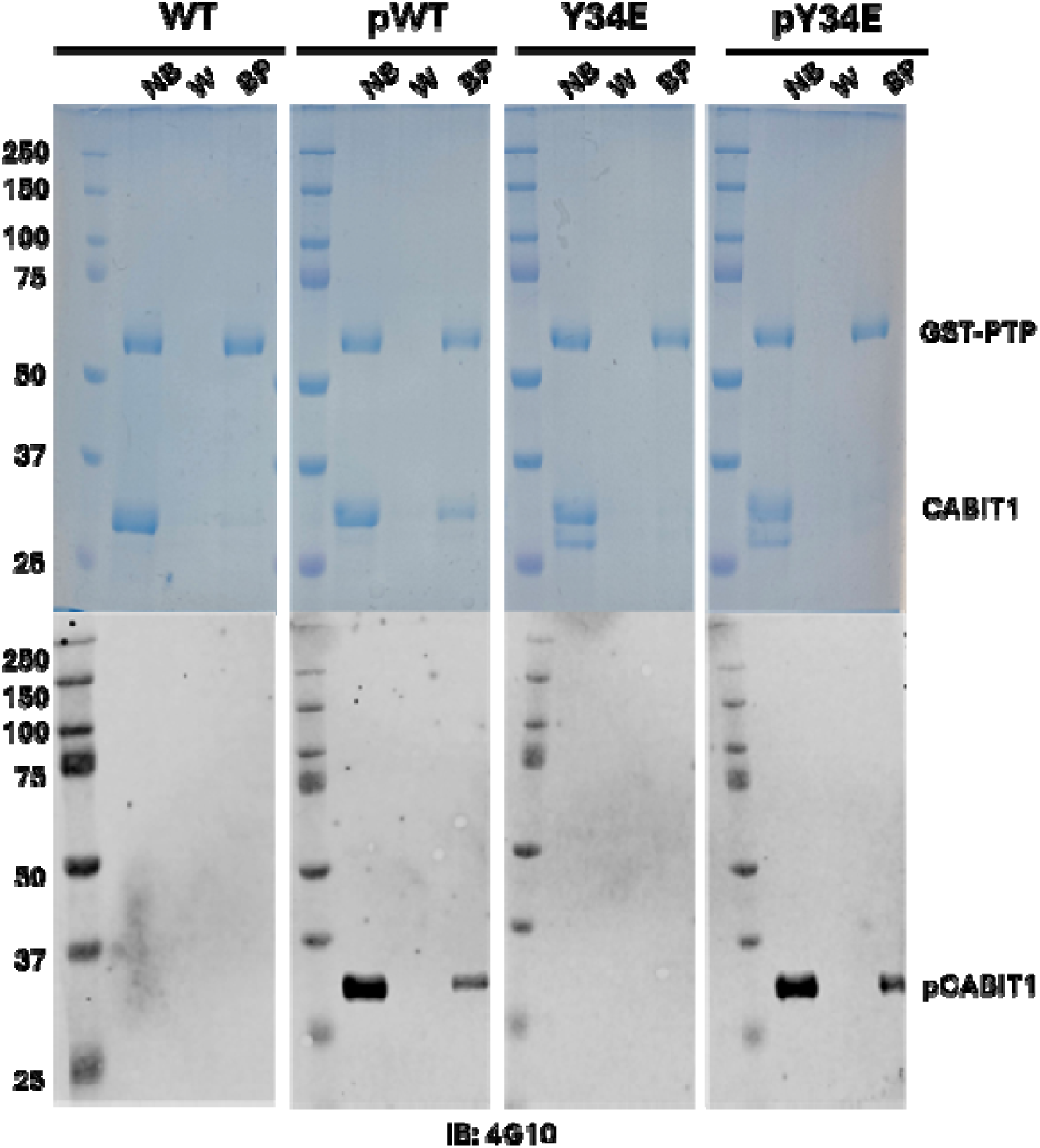
GST pulldown assays of CABIT1, pCABIT1, CABIT1 Y34E, and pCABIT1 Y34E by GST-Shp1 C453S/D419A analyzed via Coomassie blue and Western blot analysis (IB: 4G10, n=3). For both Y34 and pY34, the CABIT1 has a double band that shows up even after both affinity and size-exclusion purification.

We generated the Y34E variant of CABIT1 as a potential phosphomimetic but did not observe binding in pulldown assays with purified proteins (Figure 3). This was not surprising given the difficulty of mimicking phosphotryrosine with acidic amino acids. Phosphorylation of the Y34E variant by LCK leads to modest binding between Shp1 and CABIT1, and phospho-western blotting revealed the presence of phosphotyrosine, which must be on other sites of the domain and are likely artifacts of *in vitro* phosphorylation with isolated CABIT1 domain, as they have not been observed in cells.

### CABIT1 attenuates Shp1 inactivation by ROS

During TCR signaling, ROS are generated, which can serve as a mechanism to modulate signal transduction^25^. Shp1 is an important negative regulator of TCR signaling with a redox-sensitive Cys residue in its active site whose oxidation inactivates the phosphatase^24, 26^. A previous study showed that THEMIS can enhance and attenuate Shp1 activity, the latter involving oxidation of Shp1^14^. Given that phosphorylated THEMIS is a substrate for Shp1^13^, we hypothesized that THEMIS enhances the activity of Shp1 by shielding the active site from oxidation by ROS via a substrate-competitive mechanism. To test this, we performed phosphatase activity assays in presence of hydrogen peroxide as a source of ROS and measured the rate of oxidative inactivation of Shp1, which is irreversible under non-reducing conditions we employed. Such inactivation follows a two-step mechanism in which the first, reversible inhibition is expressed by K_i_ and the second irreversible step expressed by k_inact_. Often, experimental design limitations prevent determination of individual values and instead data are reported as k_inact_/K_i_. We determined the effect of phosphorylated and unphosphorylated CABIT1 on K_inact_/K_i_ for Shp1 inhibition by H_2_O_2_ (Figure 4A, Supplemental Figure 3). There was a statistically significant decrease (p=0.0144) in Shp1 PTP inactivation rate by H_2_O_2_ in the presence of CABIT1. Phosphorylated CABIT1 produced a more pronounced and significant decrease (p=0.0057) in Shp1 inactivation. (Figure 4B). We included Grb2, the CABIT2 domain of THEMIS, and BSA as controls as these proteins are not known to bind the Shp1 PTP domain. These proteins all decreased the rate of Shp1 inactivation but to a lesser extent than pCABIT1, demonstrating the enhanced ability of pCABIT1 to attenuate oxidative inhibition of Shp1.

**Figure 4:**
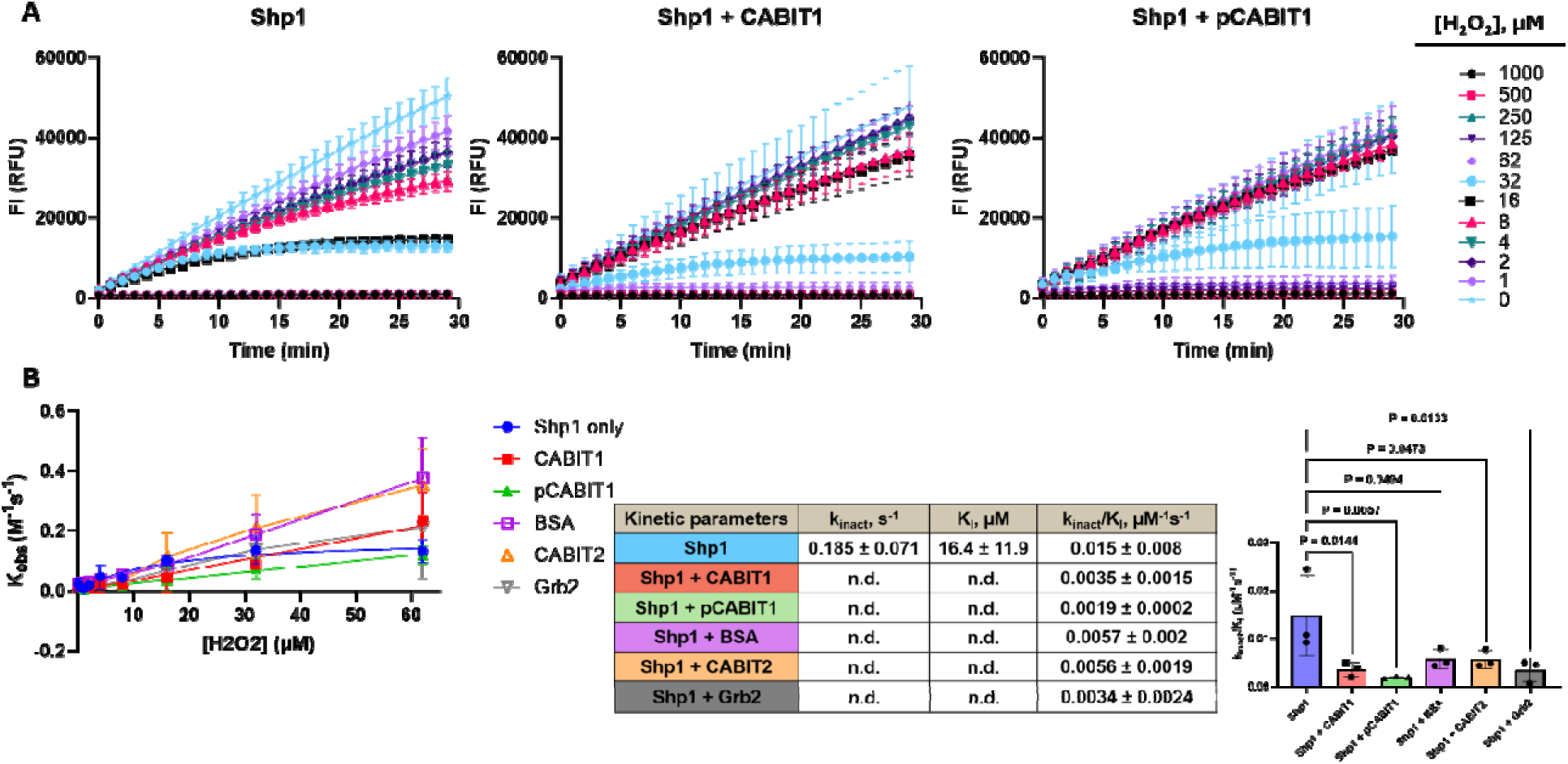
Kinetics of Shp1 oxidative inhibition. **A)** Reaction progress curves with Shp1 PTP (1 µM) and varying concentrations of H_2_O_2_. Reactions were repeated in the presence of CABIT1 (1 µM) and pCABIT1 (1 µM). Reactions were performed with 50 µM DiFMUP as substrate (n=3, mean ± SD). **B)** K_obs_ as a function of H_2_O_2_ concentration (n=3, mean ± SD). The table reports the kinetic parameters derived from both linear and non-linear regression, as described in the methods section (mean ± SD).

## Discussion

While THEMIS does not have a known enzymatic function, it has been characterized as a key component of TCR signaling through a complex with other proteins^8,10,13–14^. Cytosolic THEMIS localizes to the TCR by Grb2, which binds to THEMIS and phosphorylated LAT. This transportation allows for phosphorylation of THEMIS by LCK and subsequent SHP phosphatase binding to the CABIT1 domain^13^. However, the molecular details of the THEMIS CABI1:Shp1 interaction and how it influences Shp1 activity remains understudied. This motivated this work to determine the crystal structure of the CABIT1 domain. A portion of the CABIT1 domain was solved using molecular replacement with the AlphaFold prediction of the same domain in THEMIS and maintained a similar back bone fold (RMSD = 0.6 Å), although it’s difficult to determine if the side-chain orientation is correct due to the low-resolution data. The main difference between the structure and the prediction lies in two long β-sheets that run along the “core” subdomain of CABIT1. The other key difference is that there is no density for the loop containing Y34, which is common for dynamic regions that are post-translationally modified. Additionally, the CABIT1 domain dimerizes both in solution and in the crystal structure, but this is likely an artifact of this protein construct as similar dimerization is not observed when purifying the CABIT2 domain (Figure S1).

When aligning the CABIT2 domain structure with the CABIT1 domain prediction to each other (RMSD = 13.6 Å), we observed that both domains generally have a similar fold. These two regions were named as the “core” and the “head” subdomains respectively. The head subdomains of each CABIT domain align almost identically (Figure 2D). Meanwhile, the core subdomains have several key differences from (Figure 2C). One of the main differences lies in the overall fold of the N-terminal region of both CABIT domains, where in CABIT1 it comprises of extended loops, while CABIT2 is comprised of more ordered secondary structural folds (Figure 2C). A previous report showed that Y34 in an extended loop of CABIT1 is phosphorylated by LCK which promotes Shp1 binding (Figure 3)^13^. Another key difference between both CABIT domains is the proline rich sequence (PRS) found proximal to the C-terminus of CABIT2, whereas in CABIT1 there is only a small linker connecting the domains. The PRS found on CABIT2 has been previously characterized as the main binding region for Grb2^11–12^.

Phosphorylation of THEMIS has been shown to promote binding to Shp1, which is hypothesized to displace the autoinhibitory SH2 domains in Shp1 that occlude the active site, thereby activating the phosphatase^13^. When in the active “open” conformation, the Shp1 active site can be oxidized by ROS^14^. Our data suggests that apart from activating Shp1, it is also involved in regulating the oxidation of Shp1 by ROS. Phosphatase assay data shows that in presence of either CABIT1 or pCABIT1, Shp1 activity is preserved in the presence of H_2_O_2_ (Figure 4A and 4B as evidenced by a decrease in the inactivation coefficient (k_inact_/K_I_). A notable caveat of these experiments is the use of isolated protein domains. Shp1 exists in a basal autoinhibited conformational state that the isolated PTP domain used in these experiments lack. Thus, we expect most substrates of the Shp1 PTP domain, not only those that can activate Shp1, to provide some degree of competitive protection from oxidation in this system. Indeed, multiple proteins can slow the rate of inactivation of Shp1 PTP, though pCABIT1 provides the greatest effect (Figure 4B). Additionally, these experiments are short in duration and may not apply to all levels of cellular stress. For example, THEMIS was recently shown to promote oxidation of Shp1 in the context of chronic viral infection and T cell exhaustion via the PD-1 pathway^32^. ROS levels during chronic infection exceed those typically found in developing thymocytes and persist for longer. It is unlikely that pTHEMIS can protect Shp via a substrate- competitive mechanism under such chronic, high-ROS conditions, leading to oxidation and impaired PD-1 signaling. We therefore expect this competitive mechanism of reduced Shp1 inactivation to apply primarily to acute changes in cellular ROS.

## Author Contributions

KNT, DOS, and TSB conducted experiments. KNT and TSB wrote the manuscript. TSB supervised experiments. All authors reviewed and approved of this manuscript.

## Conflicts of Interest

No authors declare any competing conflicts of interest.

## Acknowledgements

We thank Emory University School of Medicine and the Winship Cancer Institute for startup funding and support. We thank SER-CAT at the Advanced Photon Source (APS) for their support during the APS upgrade. The Berkeley Center for Structural Biology is supported by the Howard Hughes Medical Institute, Participating Research Team members, and the National Institutes of Health, National Institute of General Medical Sciences, ALS-ENABLE grant P30 GM124169. The Advanced Light Source is a Department of Energy Office of Science User Facility under Contract No. DE-AC02-05CH11231. The Eiger detector on beamline 2.0.1 was funded under NIH grant 1S10OD032212. The Pilatus detector on beamline 5.0.1 was funded under NIH grant S10OD026941.

## Notes

### Competing Interest Statement

The authors have declared no competing interest.

### Summary of Updates

Updates to stats and results for Figure 4; added Supplemental Figure 3.

