## Supplemental figures and Table 1 for "Structural and Biochemical Analysis of the CABIT1 Domain of THEMIS"

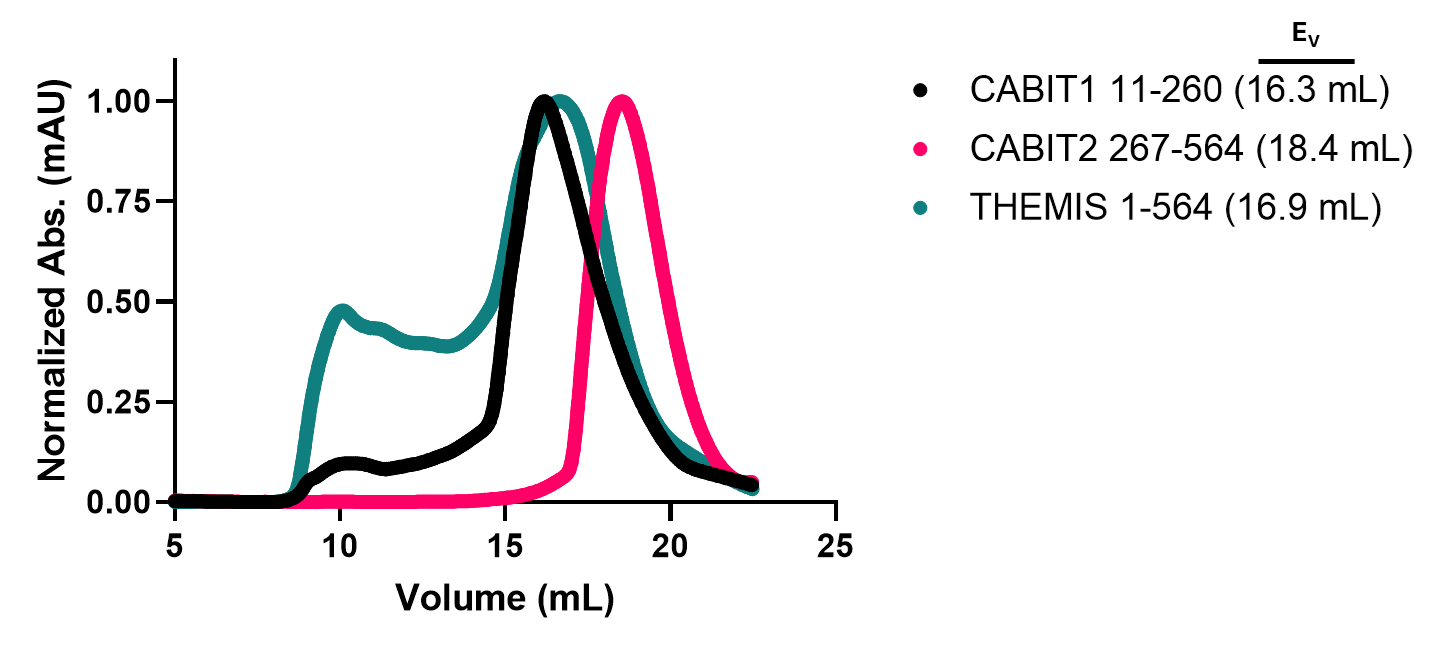

**Supplemental Figure 1:** Size-exclusion chromatograms of individual THEMIS CABIT domains and tandem CABIT domains (1-564).

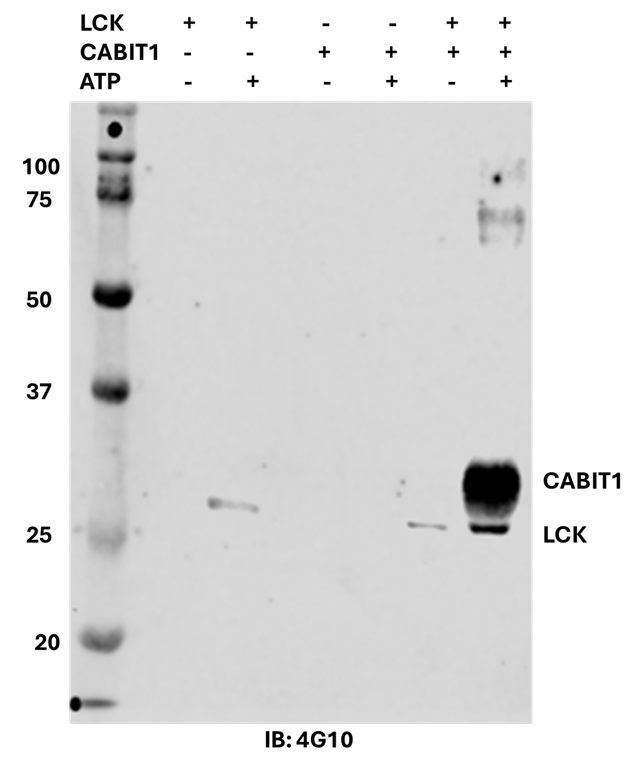

**Supplemental Figure 2:** Western blot analysis (IB: 4G10) CABIT1 domain tyrosine phosphorylation (20 µg) by LCK (1 µM) in presence and absence of 1mM ATP.

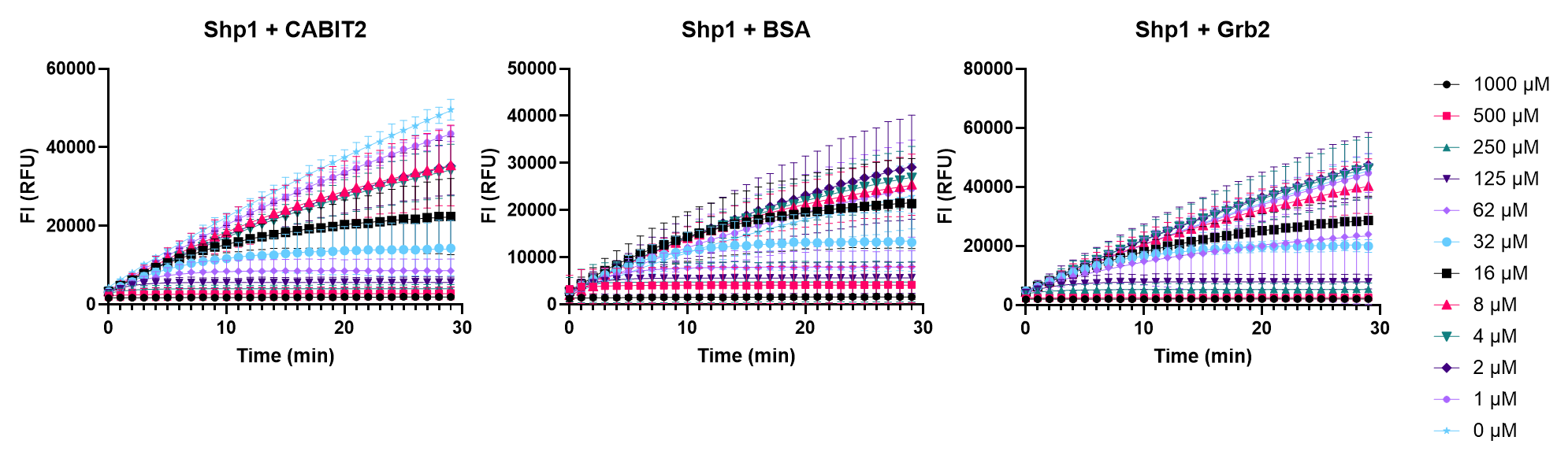
**Supplemental Figure 3:** Reaction progress curves with Shp1 PTP (1 µM) and varying concentrations of H_2_O_2_ in the presence of CABIT2 (1 µM), BSA Fraction V (1 µM), and Grb2 (1 µM). Reactions were performed with 50 µM DiFMUP as substrate (n=3, mean ± SD).

| **Construct residue boundaries** | **Bacterial Expression Results** | **Crystallization** |
| --- | --- | --- |
| **1-260** | **Mostly Insoluble** | **N/A** |
| **11-260** | **Soluble** | **Yes** |
| **15-260** | **Soluble** | **Yes** |
| **18-260** | **Soluble** | **Yes** |
| **1-268** | **Soluble** | **No** |
| **11-268** | **Soluble** | **No** |
| **18-268** | **Soluble** | **No** |
| **1-277** | **Insoluble** | **N/A** |
| **1-183** | **Insoluble** | **N/A** |
| **1-191** | **Insoluble** | **N/A** |
| **1-222** | **Insoluble** | **N/A** |
| **1-238** | **Soluble** | **N/A** |
| **1-265** | **Soluble** | **N/A** |
| **11-254** | **Soluble** | **Yes** |
| **11-256** | **Soluble** | **Yes** |
| **15-256** | **Soluble** | **Yes** |
| **15-254** | **Soluble** | **Yes** |
| **15-256** | **Soluble** | **Yes** |
| **18-254** | **Soluble** | **Yes** |
| **18-256** | **Soluble** | **Yes** |

**Supplemental Table 1:** Constructs of CABIT1 that were tested for expression and for crystallization. The constructs attempted for crystallography have had their fusion tag cleaved as described in the standard crystallization conditions, as detailed in the Methods section. N/A, not attempted.

**Table1:** Data collection and processing statistics.

|  | THEMIS CABIT1 |
| --- | --- |
| PDB Accession | 10WM |
| Wavelength (Å) | 1.00004 |
| Resolution range (Å) | 48.31 - 3.81 (3.946 - 3.81) |
| Space group | C 1 2 1 |
| Unit cel. *a, b, c (Å)*  *⍺, β, γ (°)* | 63.798 94.476 94.770  90 98.16 90 |
| Total reflections | 14977 (1484) |
| Unique reflections | 5174 (512) |
| Multiplicity | 2.9 (2.9) |
| Completeness (%) | 92.34 (87.64) |
| Mean I/σ(I) | 8.53 (0.49) |
| Wilson B-factor | 164.41 |
| R_merge_ | 0.117 (1.884) |
| R_meas_ | 0.1433 (2.313) |
| R_pim_ | 0.0812 (1.319) |
| CC_1/2_ | 0.997 (0.17) |
| Reflections used in refinement | 5074 (468) |
| Reflections used for R-free | 506 (47) |
| R-work | 0.392 (0.478) |
| R-free | 0.446 (0.486) |
| CC_work_ | 0.818 (0.302) |
| CC_free_ | 0.700 (0.408) |
| Number of non-hydrogen atoms | 2879 |
| macromolecules | 2879 |
| ligands | 0 |
| solvent | 0 |
| Protein residues | 352 |
| RMS(bonds) | 0.004 |
| RMS(angles) | 0.66 |
| Ramachandran favored (%) | 94.83 |
| Ramachandran allowed (%) | 4.31 |
| Ramachandran outliers (%) | 0.86 |
| Rotamer outliers (%) | 1.52 |
| Clashscore | 9.75 |
| Average B-factor | 234.49 |
| macromolecules | 234.49 |
